# Testing immediate dosage compensation by irradiation of heavy-ion beams to *Drosophila miranda*

**DOI:** 10.1101/2023.04.10.536214

**Authors:** Masafumi Ogawa, Kazuhide Tsuneizumi, Tomoko Abe, Masafumi Nozawa

## Abstract

Many organisms with heteromorphic sex chromosomes have a mechanism of dosage compensation (DC) in which X-linked genes are upregulated in males to mitigate dosage imbalance between sexes and between chromosomes. However, how quickly the DC is established during evolution remains elusive. In this study, irradiating the heavy-ion beams to *Drosophila miranda* that have young sex chromosomes, the so-called neo-sex chromosomes, we induced deletions on the neo-Y chromosome to mimic the situation of Y-chromosome degeneration in which functional neo-Y-linked genes were just nonfunctionalized and tested if their neo-X-linked gametologs were immediately upregulated. Since the males with the 2-Gy irradiation of iron-ion beam showed a lower fertility, we sequenced the genomes and transcriptomes of six F_1_ males derived from these males. Our pipeline identified 82 neo-Y-linked genes in which deletions were predicted in the F_1_ males. However, all but three of them had paralogs in addition to their neo-X-linked gametologs. Moreover, candidate deletions in the remaining three genes that showed one-to-one gametologous relationship with the neo-X-linked genes occurred in UTRs and did not affect the expression levels of these genes. Therefore, we were unable to directly evaluate whether DC immediately operated on the neo-X-linked genes in response to the disruption of their neo-Y-linked gametologs. Yet, our observation that the deletions occurred less frequently in one-to-one gametologs indirectly suggests that DC unlikely operated on the neo-X-linked genes immediately after the pseudogenization of their neo-Y-linked gametologs in *D. miranda*. Therefore, dosage imbalance due to deletions in the neo-Y-linked genes without paralogs may not have effectively been compensated and individuals with such deletions could have become lethal. We speculate that the neo-sex chromosomes in *D. miranda* may be too young to establish the immediate DC. Future studies on sex chromosomes with different ages will further evaluate our tentative conclusion.

---

Sex chromosomes are thought to have originated from a pair of autosomes (Vicoso 2019). Meiotic recombination between the X chromosome (X, hereafter) and the Y chromosome (Y, hereafter) is then suppressed in many cases to maintain stable sex determination, which results in massive pseudogenization of genes on the Y except for the genes involved in male determination and sexual antagonism (Charlesworth et al., 2005). Since the losses of Y-linked genes cause the dosage imbalance between sexes (i.e., one copy and two copies of X-linked genes in males and females, respectively), the X in many organisms developed the mechanism, so-called dosage compensation (DC), to mitigate such imbalance (Ohno 1967). In *Drosophila melanogaster*, for example, the protein-RNA complex named male-specific lethal (MSL complex, hereafter) globally recruits histone acetylation to the entire male X, which triggers the doubling of the expression of X-linked genes in males (Lucchesi and Kuroda 2015).

However, since many Y-linked genes are still functional at the early phase of sex chromosome differentiation, the global DC on the X-linked genes whose Y-linked gametologs are functional seems to cause over-expression of the genes. Thus, DC on the X-linked genes may need to operate more locally only when their Y-linked gametologs are nonfunctionalized in the early stage of sex chromosome evolution. In this context, the young sex chromosomes, the so-called neo-sex chromosomes, that were formed by a fusion of an autosome with an ordinary sex chromosome have been utilized to understand the early stage of sex chromosome evolution. One of such neo-sex chromosomes emerged in *D. miranda* by a fusion of the third chromosome with the Y about 1.1 Mya (Steinemann and Steinemann 1998; Bachtrog and Charlesworth 2002). Previous studies reported that the global DC via the MSL complex is already established on the neo-X chromosome (neo-X, hereafter) in *D. miranda* (Alekseyenko et al., 2013; Zhou et al., 2013), but the extent of the global DC is incomplete (Nozawa et al., 2018; Nozawa et al., 2021). In addition, these studies found that the DC on the neo-X-linked genes with the pseudogenized neo-Y-linked gametologs is greater than that on the neo-X-linked genes with the functional neo-Y-linked gametologs (Nozawa et al., 2018; Nozawa et al., 2021). This observation suggests that not only the global DC but also more localized DC (gene-by-gene DC, hereafter) operates on the neo-X by recognizing the functionality of neo-Y-linked genes in *D. miranda*, although the underlying mechanism and the immediacy of gene-by-gene DC remains unknown.

In this study, we therefore tackled how quickly such gene-by-gene DC can be established during the evolution of sex chromosomes. For this purpose, we mimicked the Y-chromosome degeneration by irradiating heavy-ion beams to *D. miranda*. The heavy-ion beam irradiation has been used as a mutagen and known to induce larger deletions than X-ray and gamma-ray (e.g., Tanaka et al., 2010). When heavy-ion (e.g., iron, argon, and carbon) beams are irradiated to the *D. miranda* males, some genomic regions may be deleted in their sperms (Fig. 1). Therefore, F_1_ males derived from a cross of the irradiated males and wildtype females may contain deletions on the neo-Y (and/or on autosomes as well but not on X or neo-X), which can be examined by genome sequencing of the F_1_ males. If a deletion is detected in a functional neo-Y-linked gene of a F_1_ male, the expression level of the neo-X-linked gametolog in the F_1_ and the wildtype males is compared to test if the DC immediately operates on the neo-X-linked gene in the F_1_ male.

**Fig. 1.**
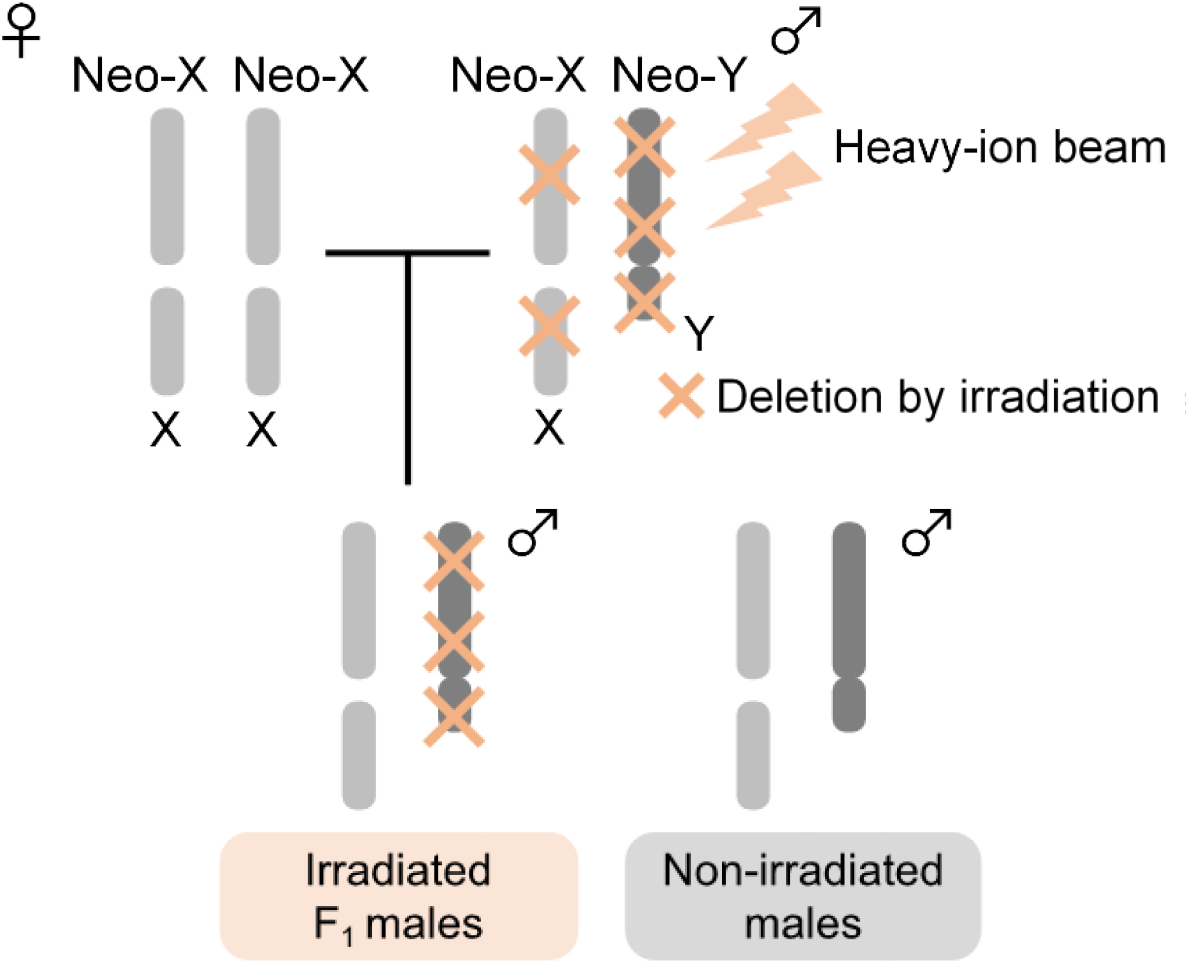
Crossing scheme to mimic Y-chromosome degeneration. Three types of heavy-ion beams (iron: Fe, argon: Ar, and carbon: C) were irradiated to *Drosophila miranda* males. The F_1_ males derived from a cross between the irradiated males and wildtype females may contain deletions on the neo-Y-linked genes. Therefore, the genomes and transcriptomes of the six F_1_ males were determined and compared with those of non-irradiated males.

First, we examined the effects of the heavy-ion beam irradiation on male fertility of *D. miranda* by using non-irradiated males as controls (Fig. 2, see also Fig. S1 for the detailed scheme of crossing experiments). The effects of the three heavy-ion beams, i.e., iron (Fe), argon (Ar), and carbon (C) ions, were separately analyzed. The 0.5-Gy irradiation of the Fe-ion beam was unlikely to severely affect male fertility (Fig. 2A, D, Table S1). In reality, the fertility of males with the 0.5-Gy irradiation after 1 day from irradiation was even greater than that without irradiation, although the variance among replicates was large. By contrast, the males with the 1-Gy and 2-Gy irradiations of the Fe-ion beam apparently showed lower fertility, particularly after 4-6 days from irradiation (Fig. 2A, D, Table S1). For the Ar- and C-ion beams, none of the irradiation levels examined (1 Gy and 2 Gy for the Ar-ion beam and 5 Gy and 10 Gy for the C-ion beam) severely affected male fertility (Fig. 2B-C, E-F, Tables S2, S3).

**Fig. 2.**
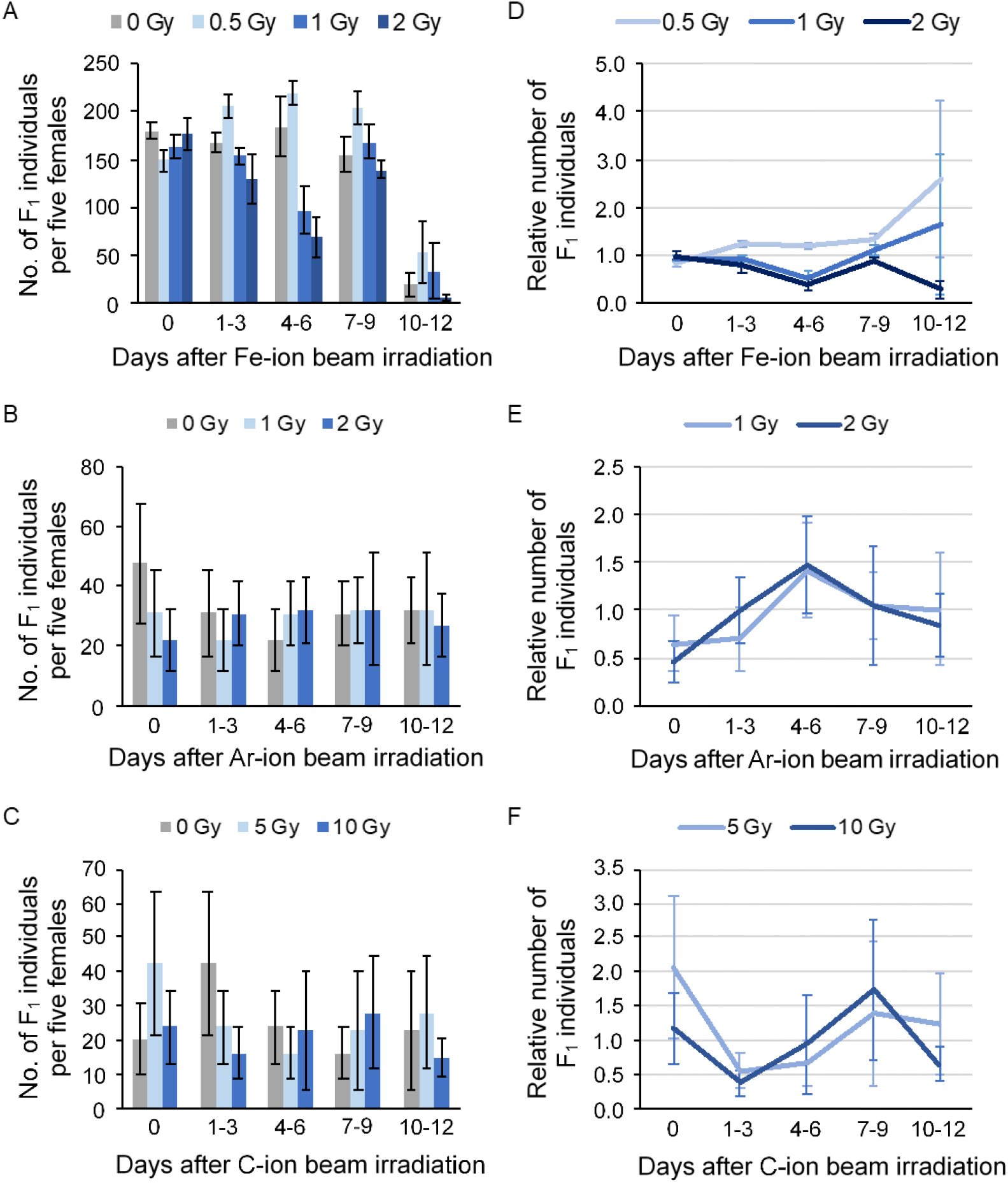
Effects of heavy-ion beam irradiation on male fertility of *Drosophila miranda*. Number of F_1_ individuals derived from the males with the irradiation of (A, D) iron (Fe)-, (B, E) argon (Ar)-, and (C, F) carbon (C)-ion beams. (A-C) Transition in the number of F_1_ individuals (offspring) after irradiation. (D-F) Transition in the number of irradiated F_1_ flies relative to that of the non-irradiated F_1_ flies. The value of 1 indicates no difference in the average number of F_1_ flies between irradiated and non-irradiated conditions. Error bars indicate the standard error among replicates (four, three, and three for the Fe-, Ar-, and C-ion beam irradiations, respectively).

Since the 2-Gy irradiation of the Fe-ion beam clearly reduced male fertility, possibly due to the deletions in the genomes of the sperms, the genomes of six F_1_ males derived from the irradiated males were sequenced to identify the deletions (Table S4). As controls, the genomes of four non-irradiated males were also sequenced. Comparing the read depth between the irradiated and the control genomes, candidate deletion regions were predicted. In short, the genomic regions that showed the read depth of zero in an irradiated male genome and at least one in all non-irradiated male genomes were regarded as deletions (See the Prediction of deletions section in Supplementary Materials and Methods for details). The results showed that the deletions on the Y/neo-Y were more frequent than those on the neo-X but less frequent compared with those on the X (Table 1). Note that any deletion on the neo-X and the X by irradiation was unexpected because these chromosomes in the F_1_ males were inherited from the non-irradiated females (Fig. 1). Therefore, our pipeline for detecting deletions produced false positives. We currently speculate that the intra-specific indel variations on the neo-X and the X could be one of the reasons for this observation. This speculation is plausible because the nucleotide polymorphism is expected to be three times more on the X than the Y at equilibrium. Indeed, the nucleotide polymorphism based on SNPs on the neo-X is much greater than that on the neo-Y (Bachtrog 2004; Nozawa et al., 2018). In other words, the neo-X (and the X as well) appears to be more prone to be affected by the indel polymorphism than the neo-Y. Nevertheless, our prediction also contained true positives. When we randomly selected seven regions in which deletions were predicted on the Y/neo-Y and conducted PCR, cloning, and Sanger sequencing, two out of the seven candidate regions indeed contained deletions at the nearby regions of our prediction (Table S5, Fig. S2, See also the Confirmation of candidate deletions section in Supplementary Materials and Methods for details). Here, it should be mentioned that repetitive sequences are accumulated on the Y/neo-Y, which made designing primers very difficult and inevitable to amplify off-targets. In this case, sequencing target regions would also be difficult even after cloning. Therefore, the accuracy of our pipeline is likely underestimated, although we should recognize that candidate deletions predicted by our pipeline certainly contains many false positives as well as false negatives.

**Table 1.**
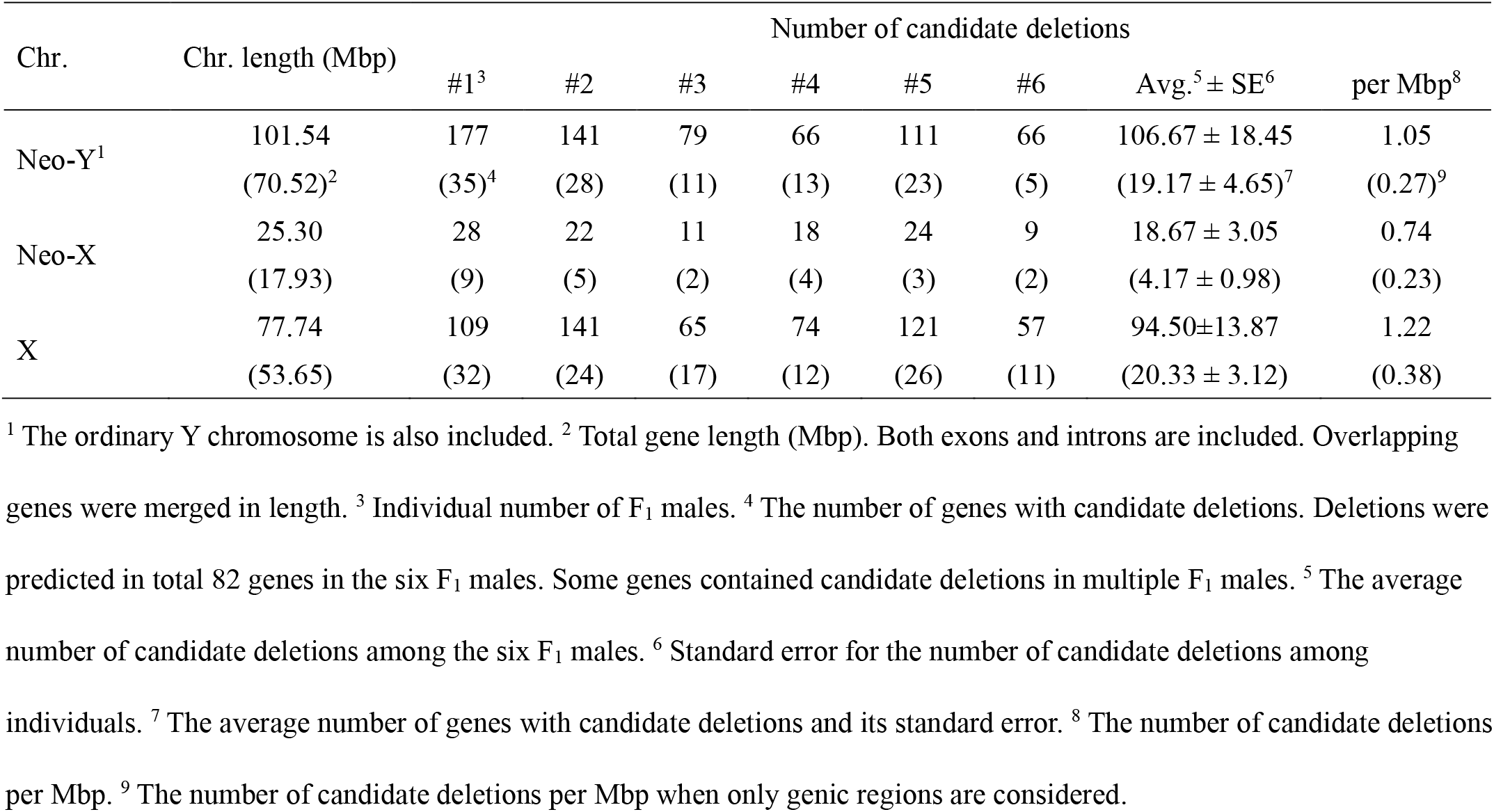
Candidate deletions by irradiation of Fe-ion beam.

The candidate deletions were distributed throughout the Y/neo-Y in the all F_1_ males (Fig. S3), indicating that the irradiation of the Fe-ion beam affected the entire chromosome more or less uniformly. The size of the largest candidate deletion on the Y/neo-Y was 1,568 bp, which was much smaller than deletions found in plants, in which the deletion size was about several hundred kb on average (Hirano et al., 2015). It is known that plant species are in general highly tolerant against irradiation. Actually, individuals with large deletions (>600 kb at maximum) were obtained by ∼400-Gy irradiation of the C-ion beam in *Arabidopsis thaliana* (Kazama et al., 2011; Kazama et al., 2017). Notably, the genome size, the number of genes, and the number of chromosomes are comparable between *Drosophila* and *Arabidopsis* (https://www.ncbi.nlm.nih.gov/genome). Therefore, further trials with greater amounts of irradiation are apparently necessary in the future to obtain large deletions effectively in *D. miranda*.

Among 9,775 genes on the Y/neo-Y, 82 genes were predicted to contain deletions in at least one of the six F_1_ males. In other words, the candidate deletions within genes were found on average every 1.24 (101.54/82) Mbp on the Y/neo-Y. It should be mentioned that after the emergence of the neo-sex chromosomes, many genes on the neo-X and the neo-Y have independently duplicated in *D. miranda* (Bachtrog et al., 2019). Therefore, the disruption of one gene among the duplicates is unlikely deleterious because paralogs would mask the effect of disruption of the gene, which would make the immediate DC dispensable, and the interpretation of the results becomes complicated. Therefore, we focused only on the genes with one copy each on the neo-X and the neo-Y (i.e., one-to-one gametologs). Then, only three one-to-one gametologs contained deletions on the neo-Y (Table 2, Fig. S3). The proportion of genes with deletions was lower for one-to-one gametologs than for other neo-Y-linked genes at marginal significance (*p* = 0.041 and 0.085 by one- and two-sided Fisher’s exact tests, respectively), suggesting the difficulty in causing deletions into one-to-one gametologs possibly due to lethality.

**Table 2.**
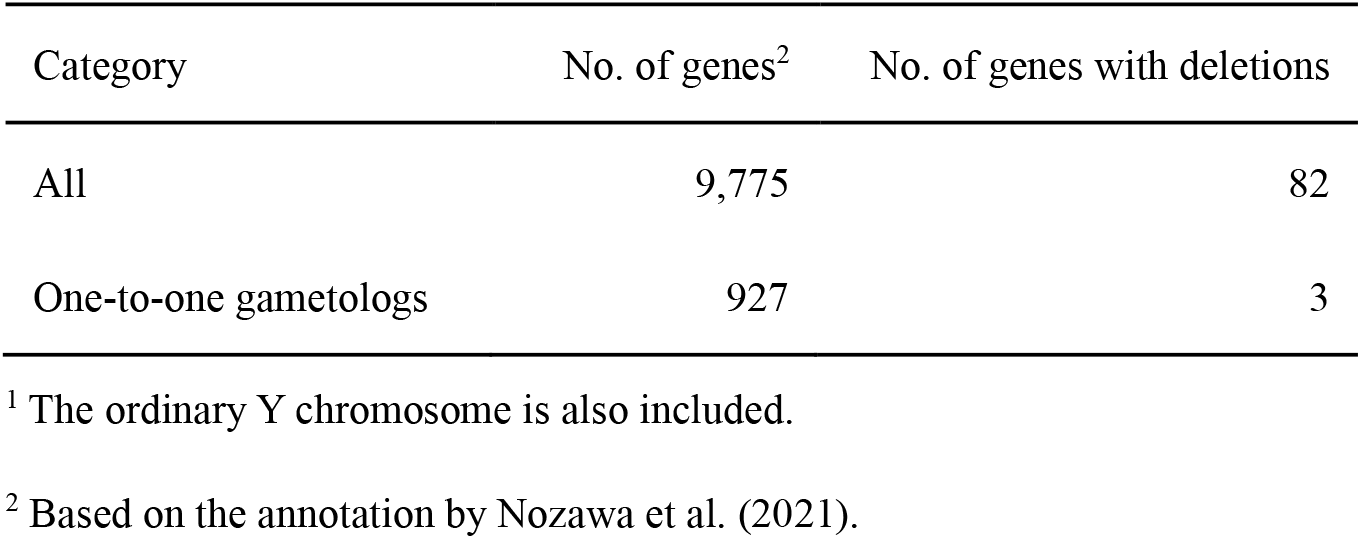
Number of genes with candidate deletions on neo-Y^1^ chromosome.

The three neo-Y-linked genes showing one-to-one gametologous relationship with the neo-X-linked genes were the homologs of *staufen* (*stau*), *back seat driver* (*bsd*), and *alicorn* (*alc*) in *D. melanogaster*. For all of them, the deletions predicted were located in UTRs (Fig. 3A) and did not change the expression level considerably in the F_1_ males, which was examined by RNA-seq (See the Gene expression analysis section in Supplementary Materials and Methods for details). Consequently, the expression levels of their neo-X-linked gametologs did not change so much (Fig. 3B-D). Therefore, we were unable to directly evaluate whether DC immediately operated in response to the disruption of the neo-Y-linked genes, because no neo-Y-linked gene was disrupted in terms of coding integrity or gene expression by the irradiation of the 2-Gy Fe-ion beam.

**Fig. 3.**
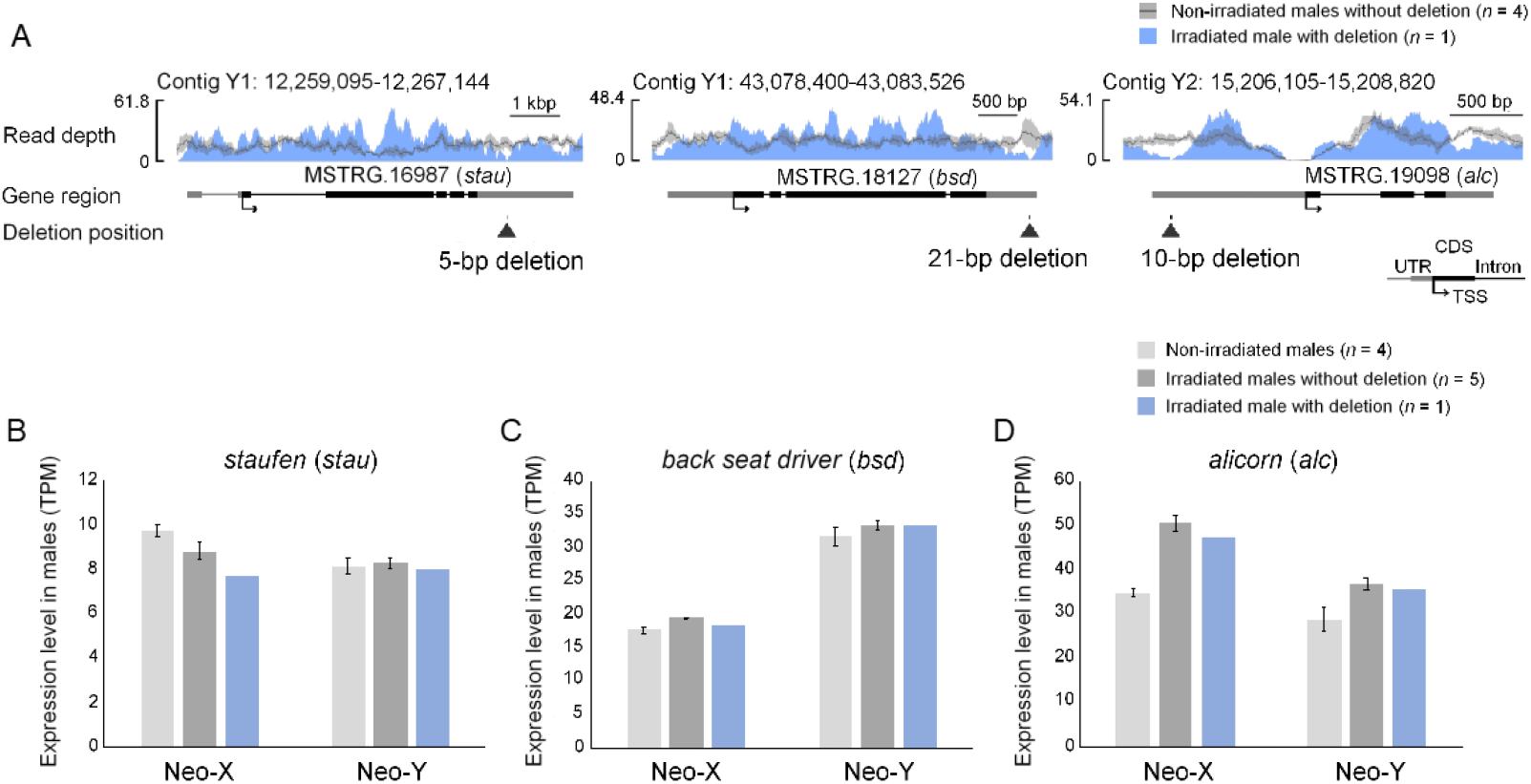
Candidate deletions in three single-copy genes on neo-Y and their effects on gene expression. (A) Positions of candidate deletions in three neo-Y-linked genes that show one-to-one gametologous relationship with their neo-X-linked genes (left: *staufen*, middle: *back seat driver*, right: *alicorn*). Positions of the deletions are indicated by arrowheads with length. Black boxes, grey boxes, and black lines indicate CDSs, UTRs, and introns, respectively. The blue track indicates the read depth of the F_1_ individual in which a deletion was detected. The grey track and dark gray line indicate the range of the read depth of the non-irradiated controls (*n* = 4) and its average, respectively. The annotation of genes was retrieved from Nozawa et al. (2021). TSS: translation start site. The plots were generated by bamCoverage (Ramirez et al., 2016) and SparK (Kurtenbach and Harbour 2019). (B-D) TPM (transcripts per million) values of (B) *staufen*, (C) *back seat driver*, and (D) *alicorn* estimated by the RNA-seq data. Neo-X: the neo-X-linked gametologs; Neo-Y: the neo-Y-linked gametologs. Light grey, dark grey, and blue bars represent the expression levels of non-irradiated males (*n* = 2), irradiated F_1_ males without deletion (*n* = 5), and an irradiated F_1_ male with deletion (*n* = 1), respectively. Error bars indicate the standard error.

Yet, the pattern that the neo-Y-linked genes showing one-to-one gametologous relationship with the neo-X-linked genes were less likely to contain deletions compared with other neo-Y-linked genes with paralogs suggests that the deletions in such single-copy genes on the neo-Y are more deleterious than those in the multi-copy genes. Note that if up-regulation of the neo-X-linked gametologs occurred immediately after the deletions in the single-copy neo-Y-linked genes, such deletions would have been dispensable and unlikely deleterious. Therefore, the deletion patterns of the neo-Y-linked genes obtained in this study may indirectly suggest that DC did not immediately operate on the neo-X in response to the pseudogenization of the neo-Y-linked genes. Yet, further experiments on gene expression are apparently needed with different irradiation conditions.

We will also need to compare our pipeline with others based on different approaches. For example, a pipeline for detecting deletions in some plants including *Arabidopsis thaliana* (Ishii et al., 2017) used the distance between the forward and reverse reads of each pair rather than the read depth adopted in this study. In this approach, deletions are detected if the distance between the forward and reverse reads of a pair on the reference genome is significantly longer than expected. It should be mentioned that this pipeline seems inappropriate to detect small deletions and was therefore unable to be applied in this study. Yet, once conditions to obtain large deletions are established, we will use this approach as well and compare it with our pipeline. It should also be mentioned that our pipeline is designed to identify the deletions on the haploid chromosomes (i.e., the X and the Y in males, see the Prediction of deletions section in Supplementary Materials and Methods for details). Therefore, deletions on autosomes were unable to be detected in this study, although deletions must have occurred on autosomes as well. We would like to improve our method to detect such heterozygous deletions on autosomes and compare the effects of deletions on autosomes and sex chromosomes in future.

In conclusion, although no neo-Y-linked single-copy genes containing deletions in coding regions were obtained, the disruption of single-copy genes on the neo-Y was less frequent than that of the multi-copy genes. This observation is consistent with the idea that the DC was not immediately established on the neo-X in response to the neo-Y degeneration. In other words, the gene-by-gene DC on the neo-X in *D. miranda* was unlikely to operate in one generation after the nonfunctionalization of neo-Y-linked genes. Since the global DC is also known to operate only partially (Nozawa et al., 2018; Nozawa et al., 2021), it is possible that the neo-sex chromosomes in *D. miranda* may be too young to establish the stringent mechanism of DC. Yet, further evaluations with different conditions of irradiation and improved pipelines of detecting deletions are apparently needed. In addition, investigating other species with different ages of sex chromosomes are also necessary to understand the relationship between the age of the sex chromosomes and the stringency of immediate DC.

We thank Yasuko Ichikawa and Yukari Kobayashi for their help on the experiments. We also thank Aya Takahashi, Reika Sato, and Takehiro K. Katoh for their comments on the manuscript. We appreciate Koichiro Tamura, Yasukazu Okada, and Yuuya Tachiki for critical discussion. This work was supported by JSPS KAKENHI Grant Numbers 21H02539 and 19K22460 to M.N. Author contributions are as follows: M.N. and M.O. designed the research. M.O., M.N., K.T., and T.A. conducted experiments. M.O. and M.N. analyzed the data. M.N. and M.O. wrote the manuscript. All authors carefully checked the manuscript and approved the research contents. All sequence data generated in this study have been submitted to the DDBJ Sequence Read Archive (DRA, https://ddbj.nig.ac.jp/search) under the accession number DRA015898.

## References

Alekseyenko, A. A., Ellison, C. E., Gorchakov, A. A., Zhou, Q., Kaiser, V. B., Toda, N., Walton, Z., Peng, S., Park, P. J., Bachtrog, D. et al. (2013) Conservation and de novo acquisition of dosage compensation on newly evolved sex chromosomes in Drosophila. Genes Dev 27, 853–858.

Bachtrog, D. (2004) Evidence that positive selection drives Y-chromosome degeneration in Drosophila miranda. Nat Genet 36, 518–522.

Bachtrog, D., and Charlesworth, B. (2002) Reduced adaptation of a non-recombining neo-Y chromosome. Nature 416, 323–326.

Bachtrog, D., Mahajan, S., and Bracewell, R. (2019) Massive gene amplification on a recently formed Drosophila Y chromosome. Nat Ecol Evol 3, 1587–1597.

Charlesworth, D., Charlesworth, B., and Marais, G. (2005) Steps in the evolution of heteromorphic sex chromosomes. Heredity 95, 118–128.

Hirano, T., Kazama, Y., Ishii, K., Ohbu, S., Shirakawa, Y, and Abe, T. (2015) Comprehensive identification of mutations induced by heavy-ion beam irradiation in Arabidopsis thaliana. Plant J 82, 93–104.

Ishii, K., Kazama, Y., Hirano, T., Hamada, M., Ono, Y., Yamada, M., and Abe, T. (2017) AMAP: A pipeline for whole-genome mutation detection in Arabidopsis thaliana. Genes Genet Syst 91, 229–233.

Kazama, Y., Hirano, T., Saito, H., Liu, Y., Ohbu, S., Hayashi, Y., and Abe, T. (2011) Characterization of highly efficient heavy-ion mutagenesis in Arabidopsis thaliana. BMC Plant Biol 11, 161.

Kazama, Y., Ishii, K., Hirano, T., Wakana, T., Yamada, M., Ohbu, S., and Abe, T. (2017) Different mutational function of low- and high-linear energy transfer heavy-ion irradiation demonstrated by whole-genome resequencing of Arabidopsis mutants. Plant J 92, 1020–1030.

Kurtenbach, S., and Harbour, J. W. (2019) SparK: A Publication-Quality NGS Visualization Tool. bioRxiv 845229.

Lucchesi, J. C., and Kuroda, M. I. (2015) Dosage compensation in Drosophila. Cold Spring Harb Perspect Biol 7.

Nozawa, M., Ikeo, K., and Gojobori, T. (2018) Gene-by-Gene or Localized Dosage Compensation on the Neo-X Chromosome in Drosophila miranda. Genome Biol Evol 10, 1875–1881.

Nozawa, M., Minakuchi, Y., Satomura, K., Kondo, S., Toyoda, A., and Tamura, K. (2021) Shared evolutionary trajectories of three independent neo-sex chromosomes in Drosophila. Genome Res 31, 2069–2079.

Ohno, S. (1967) Sex chromosomes and sex-linked genes. Springer-Verlag.

Ramirez, F., Ryan, D. P., Gruning, B., Bhardwaj, V., Kilpert, F., Richter, A. S., Heyne, S., Dundar, F., and Manke, T. (2016) deepTools2: a next generation web server for deep-sequencing data analysis. Nucleic Acids Res 44, W160–165.

Steinemann, M., and Steinemann, S. (1998) Enigma of Y chromosome degeneration: neo-Y and neo-X chromosomes of Drosophila miranda a model for sex chromosome evolution. Genetica 102–103, 409-420.

Tanaka, A., Shikazono, N., and Hase, Y. (2010) Studies on biological effects of ion beams on lethality, molecular nature of mutation, mutation rate, and spectrum of mutation phenotype for mutation breeding in higher plants. J Radiat Res 51, 223–233.

Vicoso, B. (2019) Molecular and evolutionary dynamics of animal sex-chromosome turnover. Nat Ecol Evol 3, 1632–1641.

Zhou, Q., Ellison, C. E., Kaiser, V. B., Alekseyenko, A. A., Gorchakov, A. A., and Bachtrog, D. (2013) The epigenome of evolving Drosophila neo-sex chromosomes: dosage compensation and heterochromatin formation. PLoS Biol 11, e1001711.

