## Supplementary Materials for "Testing immediate dosage compensation by irradiation of heavy-ion beams to *Drosophila miranda*"

### Supplementary Materials and methods

#### Fly strain

*Drosophila miranda* (strain 14011-0101.17) was obtained from the National Drosophila Species Stock Center (https://www.drosophilaspecies.com/) and has been maintained as a living stock at Tokyo Metropolitan University.

#### Irradiation of heavy-ion beam

Adult virgin males with 3- or 4-days after eclosion were subjected to the heavy-ion beam irradiation. Five males were put into a 15-ml tube containing ~3 ml of 50% grape juice with 1% Agar. To effectively expose the flies to the heavy-ion beam as much as possible, the medium was sloped by ~30 degrees (Fig. S1A).

Irradiation sources used in this study were iron-ion (Fe: 806 keV/µm, 0.5, 1, and 2 Gy), argon-ion (Ar: 189 keV/µm, 1 and 2 Gy), and carbon-ion (C: 30 keV/µm, 5 and 10 Gy). The ion beams were irradiated at the RI Beam Factory in RIKEN Nishina center.

For each condition, the irradiated males were crossed with five wildtype virgin females on the same day as irradiation in the standard cornmeal medium at 20°C. To examine time-course fertility of males after irradiation, on the 1st, 4th, 7th, and 10th days after irradiation, only the males were transferred to a new medium and crossed with the different virgin females (Fig. S1B). All females were discarded from a vial three days after the cross. Thus, the five vials with females crossed with males after 0, 1-3, 4-6, 7-9, and 10-12 days from irradiation were set up. All $F_1$ flies in each vial was counted until no more eclosion was observed (Tables S1-S3). To consider the variance of the experiments, we made four, three, and three replicates for the Fe-, Ar-, and C-ion beam irradiations, respectively.

#### Extraction of DNA and RNA and sequencing

Five $F_1$ adult males (from the 1-3 days vial with the condition of 2-Gy Fe-ion beam irradiation) and one $F_1$ adult male (4-6 days, 2-Gy Fe-ion) were flash-frozen in liquid nitrogen. Each fly was separately crushed and homogenized in Lysis Buffer RP1 packaged in NucleoSpin TriPrep (Takara, Shiga, Japan). Both genomic DNA and total RNA of each sample were extracted with this kit. DNA-seq libraries were constructed with NEBNext Ultra II FS DNA Library Prep Kit for Illumina (NEB, Ipswich, USA). For RNA-seq, mRNA was selected with NEBNext Poly(A) mRNA Magnetic Isolation Module (NEB), and RNA-seq libraries were constructed with NEBNext Ultra II RNA Library Prep Kit for Illumina (NEB). The 151-bp paired-end sequencing was performed on HiSeq X (Illumina, San Diego, USA) by Macrogen (Seoul, South Korea). As controls, DNA-seq for four non-irradiated males were also conducted. The data used in this study are summarized in Table S4.

#### Prediction of deletions

Deletions in an irradiated male were detected by comparing the read depth of the irradiated male with control (i.e., non-irradiated) males. In short, if there was a genomic region to which no read from the irradiated male but at least one read from all control males was mapped, the region was considered as a deletion region. For this approach, the adapter sequences and the low-quality reads (either quality <25 or length <50 at one end) were first removed from the raw data with Cutadapt v3.4 (Martin 2011) and SolexaQA++ v3.1 (Cox et al., 2010), respectively. Second, the qualified paired reads were mapped to the *D. miranda* reference genome (Nozawa et al., 2021) using bowtie2 v2.4 (Langmead and Salzberg 2012) with end-to-end and very-sensitive options. Third, the mapped reads were sorted by the genomic coordinates with the "sort" command in SAMtools v1.11 (Li et al., 2009). Fourth, PCR duplicates were removed and variants were called with the "MarkDuplicatesSpark" and "HaplotypeCaller" commands of GATK v4.2, respectively (McKenna et al., 2010). Finally, the read depth for each genomic position was counted with the "depth" command in SAMtools.

To predict deletions, we compared the read depth of each irradiated $F_1$ male with that of the control males with the custom Python script (predDeletion.py developed by M.O.), which is available upon the request to M.N. The details of this script are as follows. First, the genome sequence was converted to the array of one and zero. One denotes A, T, G, or C nucleotide, whereas zero means a non-ATGC character (e.g., N) due to incompleteness of the reference genome sequence. Second, the array of the read depth for each nucleotide position on the genome was generated from the output files of all control males by the "depth" command of SAMtools. Similarly, the array of the read depth for each position in the irradiated $F_1$ males was generated. Finally, by comparing the two arrays, the deletions in each irradiated male were predicted if the following three criteria were fulfilled: 1) the read depth is zero in the irradiated male but one or greater in all control males, 2) the region does not contain any non-ATGC characters in its surrounding 100-nucleotide positions at both ends, and 3) all of the surrounding 50-nucleotide positions at both ends show the read depths of at least one.

To identify the disrupted genes due to the deletion caused by the heavy-ion beam irradiation, the overlap between the genomic locations of the candidate deletions and genes annotated by Nozawa et al. (2021) was examined with the "intersect" command of BEDtools v2.29 (Quinlan and Hall 2010).

**Confirmation of candidate deletions**

To examine the accuracy of our pipeline to detect deletions, we selected seven candidate deletions for which primers were able to be designed in the surrounding regions (Table S5). Actually, one of the regions was partially overlapping in two $F_1$ males so that two candidate deletions were examined by a single pair of primers. First, using the DNA from a $F_1$ male with a candidate deletion, PCR was conducted by using Ex Taq (Takara) as follows: 94˚C for 2 min; 30 cycles of 94˚C for 30 sec, 60˚C for 30 sec, and 72˚C for 1 min; 72˚C for 7 min. Since the primers designed were unable to distinguish neo-Y sequences from neo-X homologous sequences, each PCR product was processed with gel extraction, ligated with the pMD20-T vector using Mighty TA-cloning Kit (Takara), and cloned into *Escherichia coli* (DH5α). Plasmid DNA in positive colonies confirmed by colony PCR was then extracted with Plasmid DNA Extraction Mini Kit (Favorgen, Ping Tung, Taiwan). Sequencing reaction was conducted with SupreDye Cycle Sequencing Kit v3.1 (AdvancedSeq, Pleasanton, USA) with either M13 forward (GTTTTCCCAGTCACGACGTT) or reverse (GGAAACAGCTATGACC ATGA) primer. Sequencing was done by using ABI3130xl Genetic Analyzer (ThermoFisher, Waltham, USA). As a control, same experiments were also conducted with wildtype males without deletions.

##### Gene expression analysis

In this study, RNA-seq was conducted for the six irradiated $F_1$ males that were also used for DNA-seq. For non-irradiated samples, the RNA-seq data derived from wildtype males generated by Nozawa et al. (2016) were used (Table S4). Gene expression analysis was performed with the same pipeline as Nozawa et al. (2021). In short, the qualified RNA-seq reads (quality ≥25 and length ≥50 at both ends) were mapped to the reference genome with STAR v2.7 (Dobin et al., 2013), and transcript per million (TPM) was calculated as the gene-expression level with RSEM v1.3 (Li and Dewey 2011).

#### References


Cox, M. P., Peterson, D. A., and Biggs, P. J. (2010) SolexaQA: At-a-glance quality assessment of Illumina second-generation sequencing data. BMC Bioinformatics **11**, 485.

Dobin, A., Davis, C. A., Schlesinger, F., Drenkow, J., Zaleski, C., Jha, S., Batut, P., Chaisson, M., and Gingeras, T. R. (2013) STAR: ultrafast universal RNA-seq aligner. Bioinformatics **29**, 15-21.

Edgar, R. C. (2004) MUSCLE: multiple sequence alignment with high accuracy and high throughput. Nucleic Acids Res **32**, 1792-1797.

Guy, L., Kultima, J. R., and Andersson, S. G. (2010) genoPlotR: comparative gene and genome visualization in R. Bioinformatics **26**, 2334-2335.

Langmead, B., and Salzberg, S. L. (2012) Fast gapped-read alignment with Bowtie 2. Nat Methods **9**, 357-359.

Li, B., and Dewey, C. N. (2011) RSEM: accurate transcript quantification from RNA-Seq data with or without a reference genome. BMC Bioinformatics **12**, 323.

Li, H., Handsaker, B., Wysoker, A., Fennell, T., Ruan, J., Homer, N., Marth, G., Abecasis, G., and Durbin, R. (2009) The Sequence Alignment/Map format and SAMtools. Bioinformatics **25**, 2078-2079.

Martin, M. (2011) Cutadapt Removes Adapter Sequences from High-Throughput Sequencing Reads. EMBnetJournal **17**, 10-12.

McKenna, A., Hanna, M., Banks, E., Sivachenko, A., Cibulskis, K., Kernytsky, A., Garimella, K., Altshuler, D., Gabriel, S., Daly, M., et al. (2010) The Genome Analysis Toolkit: a MapReduce framework for analyzing next-generation DNA sequencing data. Genome Res **20**, 1297-1303.

Nozawa, M., Minakuchi, Y., Satomura, K., Kondo, S., Toyoda, A., and Tamura, K. (2021) Shared evolutionary trajectories of three independent neo-sex chromosomes in *Drosophila*. Genome Res **31**, 2069-2079.

Nozawa, M., Onizuka, K., Fujimi, M., Ikeo, K., and Gojobori, T. (2016) Accelerated pseudogenization on the neo-X chromosome in Drosophila miranda. Nat Commun **7**, 13659.

Quinlan, A. R., and Hall, I. M. (2010) BEDTools: a flexible suite of utilities for comparing genomic features. Bioinformatics **26**, 841-842.

Tamura, K., Stecher, G., and Kumar, S. (2021) MEGA11: Molecular Evolutionary Genetics Analysis Version 11. Mol Biol Evol **38**, 3022-3027.

Table S1. Number of $F_1$ flies[1] derived from males with iron-ion beam irradiation.

| | 0 Gy | | | | | | 0.5 Gy | | | | | | 1 Gy | | | | | | 2 Gy | | | | | |
|---|---|---|---|---|---|---|---|---|---|---|---|---|---|---|---|---|---|---|---|---|---|---|---|---|
| Days | Rep1 | Rep2 | Rep3 | Rep4 | Avg.[2] | SE[3] | Rep1 | Rep2 | Rep3 | Rep4 | Avg. | SE | Rep1 | Rep2 | Rep3 | Rep4 | Avg. | SE | Rep1 | Rep2 | Rep3 | Rep4 | Avg. | SE |
| 0 | 158 | 174 | 199 | 189 | 180.0 | 9.0 | 170 | 166 | 129 | 131 | 149.0 | 11.0 | 147 | 154 | 200 | 154 | 163.8 | 12.2 | 187 | 210 | 186 | 128 | 177.8 | 17.5 |
| 1-3 | 166 | 190 | 143 | 172 | 167.8 | 9.7 | 228 | 216 | 213 | 171 | 207.0 | 12.4 | 166 | 138 | 174 | 137 | 153.8 | 9.5 | 174 | 75 | 96 | 172 | 129.3 | 25.6 |
| 4-6 | 267 | 168 | 190 | 114 | 184.8 | 31.7 | 258 | 209 | 205 | 211 | 220.8 | 12.5 | 77 | 60 | 83 | 168 | 97.0 | 24.2 | 82 | 29 | 123 | 42 | 68.9 | 21.2 |
| 7-9 | 204 | 126 | 160 | 131 | 155.3 | 17.9 | 179 | 172 | 249 | 217 | 204.3 | 17.9 | 204 | 160 | 188 | 125 | 169.3 | 17.3 | 152 | 158 | 118 | 130 | 139.5 | 9.4 |
| 10-12 | 2 | 0 | 53 | 25 | 20.0 | 12.4 | 2 | 138 | 0 | 68 | 52.0 | 32.7 | 120 | 11 | 1 | 0 | 33.0 | 29.1 | 0 | 15 | 0 | 8 | 5.8 | 3.6 |

[1] The total number of $F_1$ flies from five females was shown.

[2] Average among the replicates.

[3] Standard error among the replicates.

Table S2. Number of $F_1$ flies[1] derived from males with argon-ion beam irradiation.

| | 0 Gy | | | | | 1 Gy | | | | | 2 Gy | | | | |
|---|---|---|---|---|---|---|---|---|---|---|---|---|---|---|---|
| Days | Rep1 | Rep2 | Rep3 | Avg.[2] | SE[3] | Rep1 | Rep2 | Rep3 | Avg. | SE | Rep1 | Rep2 | Rep3 | Avg. | SE |
| 0 | 98 | 52 | 41 | 47.8 | 20.1 | 38 | 67 | 19 | 31.0 | 14.3 | 14 | 24 | 49 | 21.8 | 10.3 |
| 1-3 | 38 | 67 | 19 | 31.0 | 14.3 | 14 | 24 | 49 | 21.8 | 10.3 | 39 | 51 | 33 | 30.8 | 10.9 |
| 4-6 | 14 | 24 | 49 | 21.8 | 10.3 | 39 | 51 | 33 | 30.8 | 10.9 | 39 | 40 | 49 | 32.0 | 10.9 |
| 7-9 | 39 | 51 | 33 | 30.8 | 10.9 | 39 | 40 | 49 | 32.0 | 10.9 | 87 | 20 | 22 | 32.3 | 18.9 |
| 10-12 | 39 | 40 | 49 | 32.0 | 10.9 | 87 | 20 | 22 | 32.3 | 18.9 | 38 | 23 | 48 | 27.3 | 10.4 |

[1] The total number of $F_1$ flies from five females was shown.

[2] Average among the replicates.

[3] Standard error among the replicates.

Table S3. Number of $F_1$ flies[1] derived from males with carbon-ion beam irradiation.

| | 0 Gy | | | | | 5 Gy | | | | | 10 Gy | | | | |
|---|---|---|---|---|---|---|---|---|---|---|---|---|---|---|---|
| Days | Rep1 | Rep2 | Rep3 | Avg.[2] | SE[3] | Rep1 | Rep2 | Rep3 | Avg. | SE | Rep1 | Rep2 | Rep3 | Avg. | SE |
| 0 | 31 | 7 | 44 | 20.5 | 10.3 | 83 | 13 | 74 | 42.5 | 21.0 | 49 | 15 | 32 | 24.0 | 10.6 |
| 1-3 | 83 | 13 | 74 | 42.5 | 21.0 | 49 | 15 | 32 | 24.0 | 10.6 | 8 | 34 | 23 | 16.3 | 7.6 |
| 4-6 | 49 | 15 | 32 | 24.0 | 10.6 | 8 | 34 | 23 | 16.3 | 7.6 | 0 | 73 | 18 | 22.8 | 17.3 |
| 7-9 | 8 | 34 | 23 | 16.3 | 7.6 | 0 | 73 | 18 | 22.8 | 17.3 | 18 | 76 | 19 | 28.3 | 16.5 |
| 10-12 | 0 | 73 | 18 | 22.8 | 17.3 | 18 | 76 | 19 | 28.3 | 16.5 | 14 | 28 | 18 | 15.0 | 5.8 |

[1] The total number of $F_1$ flies from five females was shown.

[2] Average among the replicates.

[3] Standard error among the replicates.

Table S4. Summary of the DNA-seq and RNA-seq data used in this study.

| Sample | Treatment | Individual[1] | Sequencing | Platform | Layout/length (nt) | Accession | Reference |
|---|---|---|---|---|---|---|---|
| #1 | Fe 2 Gy, 1-3 days | 1 | DNA-seq | HiSeq X | Paired/151 | DRR452318 | This study |
| | | | RNA-seq | HiSeq X | Paired/151 | DRR452328 | This study |
| #2 | Fe 2 Gy, 1-3 days | 1 | DNA-seq | HiSeq X | Paired/151 | DRR452319 | This study |
| | | | RNA-seq | HiSeq X | Paired/151 | DRR452329 | This study |
| #3 | Fe 2 Gy, 1-3 days | 1 | DNA-seq | HiSeq X | Paired/151 | DRR452320 | This study |
| | | | RNA-seq | HiSeq X | Paired/151 | DRR452330 | This study |
| #4 | Fe 2 Gy, 1-3 days | 1 | DNA-seq | HiSeq X | Paired/151 | DRR452321 | This study |
| | | | RNA-seq | HiSeq X | Paired/151 | DRR452331 | This study |
| #5 | Fe 2 Gy, 1-3 days | 1 | DNA-seq | HiSeq X | Paired/151 | DRR452322 | This study |
| | | | RNA-seq | HiSeq X | Paired/151 | DRR452332 | This study |
| #6 | Fe 2 Gy, 4-6 days | 1 | DNA-seq | HiSeq X | Paired/151 | DRR452323 | This study |
| | | | RNA-seq | HiSeq X | Paired/151 | DRR452333 | This study |
| Control 1 | non-irradiated | 1 | DNA-seq | HiSeq X | Paired/151 | DRR452324 | This study |
| Control 2 | non-irradiated | 1 | DNA-seq | HiSeq X | Paired/151 | DRR452325 | This study |
| Control 3 | non-irradiated | 1 | DNA-seq | HiSeq X | Paired/151 | DRR452326 | This study |
| Control 4 | non-irradiated | 1 | DNA-seq | HiSeq X | Paired/151 | DRR452327 | This study |
| Control 5 | non-irradiated | 5 | RNA-seq | HiSeq 2000 | Paired/101 | DRR055198 | Nozawa et al. (2016) |
| Control 6 | non-irradiated | 5 | RNA-seq | HiSeq 2000 | Paired/101 | DRR055199 | Nozawa et al. (2016) |

[1] The number of individuals in each sample.

Table S5. Primers used for Sanger sequencing to examine the accuracy of our pipeline for predicting deletions.

| Region | Forward primer | Reverse primer | Sequenced region in the reference genome | Individual with a predicted deletion |
|---|---|---|---|---|
| 1 | GTCCACTGTCCAGACCACCT | CGACCAGGATTCGGTTTTGT | Contig_Y2: 4,171,669-4,172,398 | #4 (41 bp)[2] |
| 2 | CGATGCGAAAGAAGTTGTCA | ATTTCGAGTCAAACGGATCG | Contig_Y1: 32,661,592-32,662,458 | #3 (26 bp) |
| 3 | TGGGTGTACCCTGTCTCTCC | CGCTTACGAATTGCACAAGA | Contig_Y1: 26,752,705-26,753,347 | #2 (50 bp) |
| 4 | CCCTGAACAAAGGAAAACCA | ACATTTCACCTCACCCTTGC | Contig_Y1: 20,012,930- 20,013,626 | #2 (18 bp) |
| 5 | CTGGGAAACAAGCCAAGTGT | AGTTGGTCCAAGTCCACACC | Contig_Y1: 24,538,547- 24,539,012 | #2 (2 bp) |
| 6[1] | AACGATAATTTGCCGTGGAG | CATTCGCTGTCGAATCTACC | Contig_Y2: 36,092,745- 36,093,482 | #1 (140 bp) #5 (139 bp) |

[1] Two individuals (#1 and #5) were predicted to have deletions in an overlapping region. Therefore, seven candidate deletions were examined by Sanger sequencing in total.

[2] Size of candidate deletions.

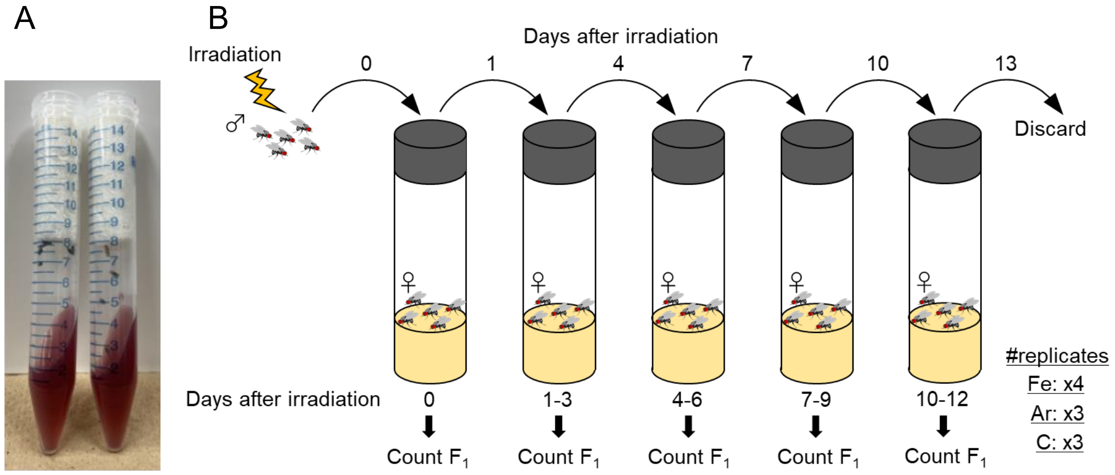


Figure S1. Experimental procedure in this study. (A) Vials used for the irradiation of heavy-ion beams. Five males were placed into each vial for irradiation. (B) Crossing scheme to obtain $F_1$ flies derived from a cross between irradiated males and wildtype females. See also the Irradiation of heavy-ion beam section in Supplementary Materials and Methods for details.

**Sample #1**

[Predicted] Contig_Y2: 36,093,216-36,093,355

[Actual] Contig_Y2: 36,093,251-36,093,257 and 36,093,276-36,093,350

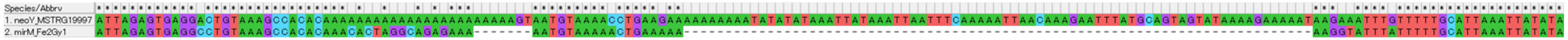


**Sample #5**

[Predicted] Contig_Y2: 36,093,197-36,093,335

[Actual] Contig_Y2: 36,092,958-36,093,013

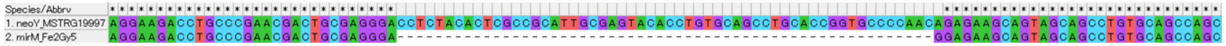


[Actual] Contig_Y2: 36,093,251-36,093,257 and 36,093,276-36,093,350

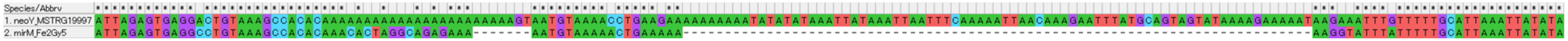


Figure S2. Confirmation of candidate deletions (region 6 in Table S5) by Sanger sequencing. “Predicted” indicated the candidate deletion regions predicted by our pipeline based on the NGS data, whereas “Actual” means the actual deletion regions determined by Sanger sequencing. Sequence alignment was conducted by Muscle (Edgar 2004) implemented in MEGA11 (Tamura et al., 2021). In each diagram, the upper and lower sequences are derived from control (non-irradiated) and irradiated males, respectively.

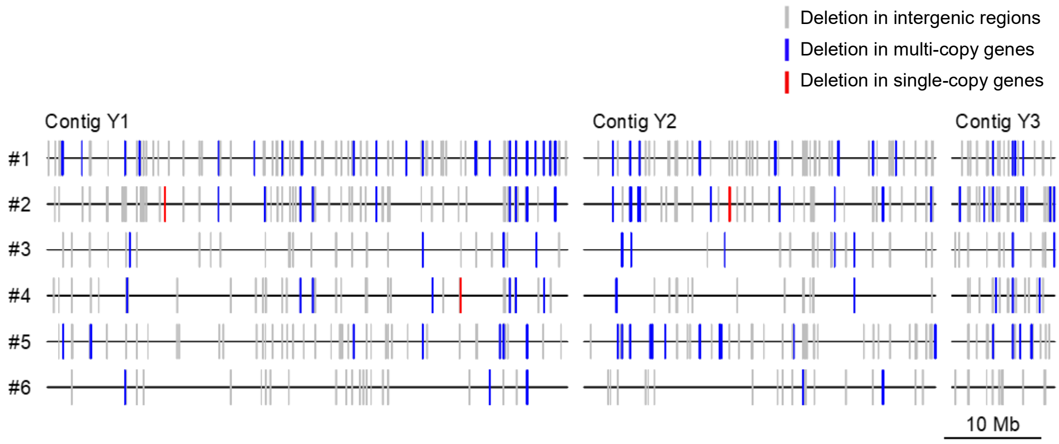


Figure S3. Distributions of candidate deletions Y/neo-Y chromosomes in six $F_1$ males derived from *Drosophila miranda* males subjected to 2-Gy Fe-ion beam irradiation. Grey, blue, and red lines indicate the candidate deletions in intergenic regions, multi-copy genes (either exons or introns), and single-copy genes (i.e., one-to-one gametologs, either exons or introns), respectively. Contigs Y1, Y2, and Y3 are the top three longest scaffolds of the Y/neo-Y in *D. miranda*. The scale-bar is shown below the diagram. The plot was generated with genoPlotR (Guy et al., 2010).
